# *In vitro* Targeting of Transcription Factors to Control the Cytokine Release Syndrome in COVID-19

**DOI:** 10.1101/2020.12.29.424728

**Authors:** Clarissa S. Santoso, Zhaorong Li, Jaice T. Rottenberg, Xing Liu, Vivian X. Shen, Juan I. Fuxman Bass

**Affiliations:** Department of Biology, Boston University, Boston, MA 02215, USA; Bioinformatics Program, Boston University, Boston, MA 02215, USA

**Author notes:** **Corresponding author:** Juan I. Fuxman Bass, Boston University, 5 Cummington Mall, Boston, MA 02215.

**Keywords:** COVID-19, cytokine release syndrome, cytokine storm, drug repurposing, transcriptional regulators

## Abstract

Treatment of the cytokine release syndrome (CRS) has become an important part of rescuing hospitalized COVID-19 patients. Here, we systematically explored the transcriptional regulators of inflammatory cytokines involved in the COVID-19 CRS to identify candidate transcription factors (TFs) for therapeutic targeting using approved drugs. We integrated a resource of TF-cytokine gene interactions with single-cell RNA-seq expression data from bronchoalveolar lavage fluid cells of COVID-19 patients. We found 581 significantly correlated interactions, between 95 TFs and 16 cytokines upregulated in the COVID-19 patients, that may contribute to pathogenesis of the disease. Among these, we identified 19 TFs that are targets of FDA approved drugs. We investigated the potential therapeutic effect of 10 drugs and 25 drug combinations on inflammatory cytokine production in peripheral blood mononuclear cells, which revealed two drugs that inhibited cytokine production and numerous combinations that show synergistic efficacy in downregulating cytokine production. Further studies of these candidate repurposable drugs could lead to a therapeutic regimen to treat the CRS in COVID-19 patients.

## Introduction

Coronavirus Disease-2019 (COVID-19), caused by the SARS-CoV-2 betacoronavirus strain, has led to over 80 million confirmed cases and 1.7 million deaths worldwide, since its first reported case in December 2019 (1). Most COVID-19 cases are either asymptomatic or cause only mild disease (2). However, a considerable number of patients develop severe respiratory illnesses manifested in fever and pneumonia, leading to acute respiratory distress syndrome (ARDS) and cytokine release syndrome (CRS) (3). CRS is an acute systemic inflammatory response characterized by the rapid and excessive release of inflammatory cytokines. Uncontrolled CRS results in systemic hyperinflammation and can lead to life-threatening multi-organ failure (3).

There is an urgent need for therapies to treat the CRS in COVID-19 patients. While government agencies and private companies have accelerated procedures to develop and distribute COVID-19 vaccines, it will take a year or longer for the population to be vaccinated. Additionally, a significant portion of the population may not get vaccinated due to reduced compliance and limited access to vaccines, or may not mount a proper protective response (e.g., immunodeficient patients). Furthermore, whether the vaccines generate a long-lasting protective response in all patients is still unknown. Since drug development and approval may take years, drug repurposing of already approved drugs is an efficient approach to identify alternative therapeutic options. At present, three repurposed drugs, remdesivir, dexamethasone, and baricitinib (in combination with remdesivir), have been found to benefit COVID-19 patients in large, controlled, randomized, clinical trials (4–6). Dexamethasone, which acts as an agonist of the glucocorticoid receptor (GR, also known as NR3C1) transcription factor (TF), is an anti-inflammatory corticosteroid (7). Indeed, corticosteroids have been shown to suppress CRS (8), and NR3C1 has been shown to transcriptionally downregulate many inflammatory cytokines overexpressed in COVID-19 patients, such as CCL2, IL1B, and IL6 (9, 10). Baricitinib, a non-steroidal anti-inflammatory drug, acts as a janus kinase (JAK) inhibitor (4). The JAK-signal transducers and activators of transcription (STAT) signaling pathway leads to the transcription of inflammatory cytokines, thus inhibition of the JAK-STAT signaling pathway decreases the production of inflammatory cytokines. Despite the efficacy of these drugs in reducing COVID-19 mortality, the effect-size is modest, suggesting the need for additional drugs or combinations to treat the CRS in COVID-19 patients. Although antibodies are well-proven strategies to block cytokine activity, approved antibodies are available for only nine cytokines (DrugBank, (11)), specifically TNF and various interleukins (ILs). However, the COVID-19 CRS primarily manifests in overproduction of chemokines (i.e. CCLs and CXCLs)(12, 13). Thus, as cytokines are highly transcriptionally regulated, there is great potential in exploring other transcriptional regulators of inflammatory cytokines involved in the COVID-19 CRS that can be targeted with approved drugs.

Here, we systematically studied the transcriptional regulators of inflammatory cytokines involved in the COVID-19 CRS to identify candidate TFs for therapeutic targeting using approved drugs. We integrated a resource of empirically identified TF-cytokine gene interactions with single-cell RNA-seq (scRNA-seq) expression data from COVID-19 patients to reveal correlated TF-cytokine gene interactions that may contribute to pathogenesis of the disease. We identified candidate TFs that could be targeted using approved drugs and investigated the potential therapeutic effect of 10 drugs on the expression of cytokines upregulated in COVID-19 patients. We also assayed 25 drug combinations and found numerous combinations that show promising synergistic efficacy in downregulating the expression of inflammatory cytokines. In summary, the present study provides a network-based approach focusing on the transcriptional regulators of inflammatory cytokines to identify candidate repurposable drugs to treat the COVID-19 CRS.

## Results

### Delineation of a COVID-19 cytokine gene regulatory network

We hypothesized that transcriptional regulators whose expressions are significantly correlated with the expression of cytokines upregulated in COVID-19 patients may play a role in the pathogenesis of the COVID-19 CRS. To identify TF-cytokine pairs correlated in expression, we integrated a published resource of 2,260 empirically tested TF-cytokine gene interactions (CytReg v2)(14) with publicly available scRNA-seq data of bronchoalveolar lavage fluid (BALF) cells from nine COVID-19 patients (GSE145926)(12) and three healthy controls (GSE145926 and GSE128033)(12, 15). Unsupervised clustering analysis of the scRNA-seq data revealed distinct clusters of ciliated epithelial cells, secretory epithelial cells, natural killer cells, neutrophils, macrophages, myeloid dendritic cells, plasmacytoid dendritic cells, CD4 T cells, CD8 T cells, B cells, and plasma cells, identified by signature genes (Supplementary Figure 1A-B). For each cell type, we identified cytokines that were significantly (Padj<0.05) upregulated in the COVID-19 patients compared to healthy controls (Supplementary Table S1), and then determined the TFs in CytReg v2 reported to functionally regulate or bind to the transcriptional control regions of these cytokines. To prioritize TFs that may have a role in the pathogenesis of the COVID-19 CRS, we generated gene regulatory networks for TF-cytokine interactions that are significantly correlated across single cells in each cell type (Supplementary Table S1) in the COVID-19 patient BALF samples.

In total, we identified 581 significantly correlated interactions between 95 TFs and 16 cytokines upregulated in the COVID-19 patients. Strikingly, 567 (97.6%) interactions displayed a positive correlation, suggesting that the cytokine upregulation is primarily mediated through activation by transcriptional activators rather than through de-repression by transcriptional repressors. The transcriptional activation could be a result of activated signaling pathways impinging on the TFs or increased TF expression. Indeed, 89 (93.7%) TFs were significantly upregulated in at least one cell type in the COVID-19 patients, many of which are known to be activated by signaling pathways in inflammation. Consistent with this, TFs in 336 of 395 (85.1%) positively correlated interactions that have a regulatory function reported in CytReg v2 were reported to activate expression of the target cytokine gene in various inflammatory contexts (14). This provides evidence that TFs displaying a positive correlation in the COVID-19 cytokine gene regulatory network can functionally activate expression of the target cytokine gene.

TFs that have widespread interactions across many cell types likely play important roles in regulating the COVID-19 CRS. Notably, IRF2, IRF7, and STAT1, were upregulated and positively correlated with multiple cytokines in all cell types. IRFs and STATs play prominent roles in viral infection by regulating interferon (IFN) production and response pathways and potentiating the expression of antiviral genes including inflammatory cytokine genes (16, 17). Indeed, dysregulation of the IFN pathways either by inborn errors or the generation of autoantibodies against type I IFNs has been associated with COVID-19 severity (18, 19). However, the robust interferon response observed in many severe COVID-19 patients also likely contributes to the CRS (13, 20, 21). Thus, the upregulation of IRF2, IRF7, and STAT, in all cell types may be driving the amplification of IFN response pathways and thereby the overproduction of inflammatory cytokines that contribute to COVID-19 CRS pathogenesis. Consistent with this, inhibiting JAK-STAT signaling with baricitinib significantly improved recovery time and survival rates among patients with severe COVID-19, likely by suppressing the CRS (4, 22).

We next focused on TF hubs, TFs that interact with many overexpressed cytokine genes, since they likely play important roles in COVID-19 CRS pathogenesis. We found that TFs with the highest number of positively correlated interactions were well-known pathogen-activated transcriptional activators, such as REL (29 interactions), STAT1 (29 interactions), IRF7 (28 interactions), and NFKB1 (27 interactions). Additionally, we found that NF-κB family members regulated the most number of unique cytokines (*e.g*., REL - 9 cytokines, RELA - 9 cytokines, and NFKB1 - 8 cytokines). This is consistent with NF-κB being a potent inducer of cytokine production and NF-κB hyperactivation being directly implicated in the CRS observed in severe COVID-19 patients (23). Collectively, these findings support that targeting NF-κB may have therapeutic benefits in controlling the CRS in COVID-19 patients.

Drug repurposing offers a viable therapeutic approach that can significantly shorten the time to deliver effective treatments to COVID-19 patients. We identified 19 TFs in the networks that are targets of FDA approved drugs (Figure 1 and Supplementary Table). Of these, NFKB1, RELA, JUN, FOS, and HIF1A, displayed the highest number of positively correlated interactions and interacted with the most number of unique cytokines. Similar to NF-κB TFs, AP-1 TFs (FOS and JUN) are critical regulators of inflammatory cytokine genes such as CCL2 and IL6 (24, 25), which are expressed at high levels in COVID-19 patients (26–28). Interestingly, CCL2 and IL6 can also induce expression of AP-1 genes and regulate activation of AP-1 proteins (29–31). Therefore, targeting AP-1 has the potential to block these positive feedback loops in addition to limiting the expression of multiple inflammatory cytokines overexpressed in COVID-19 patients.

**Figure 1.**
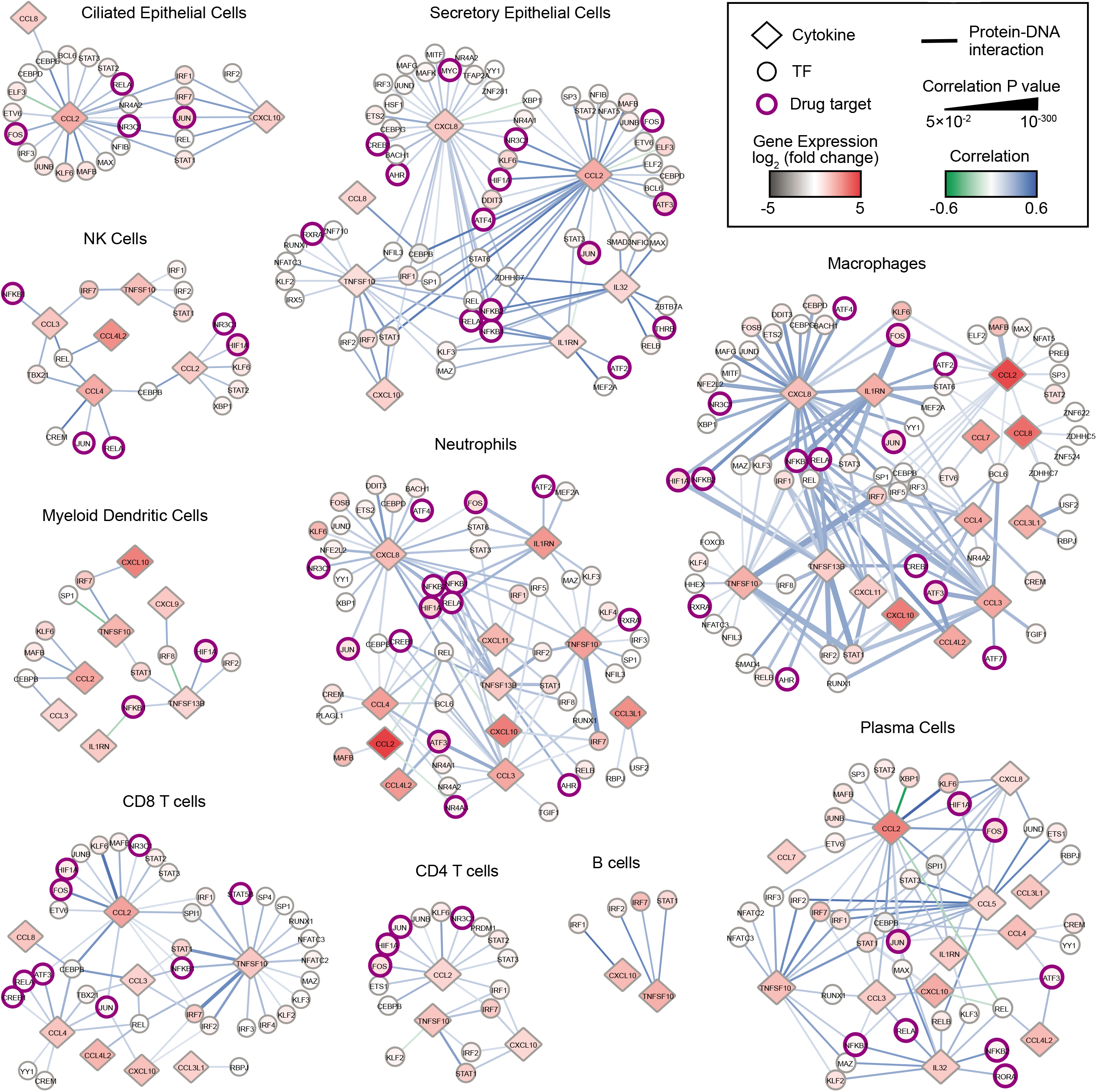
COVID-19 cytokine gene regulatory network. Immune cell sub-networks mapping 581 TF-cytokine gene interactions between 95 TFs and 16 cytokine genes upregulated in BALFs of COVID-19 patients. Networks were restricted to interactions that are significantly correlated (Padj<0.05) with a Pearson correlation coefficient >0.1 or <−0.1 in the respective immune cell subtype. Diamonds represent cytokines and circles represent TFs. TFs that are targets of FDA approved drugs are indicated in purple circles. The node color denotes the differential gene expression of TFs and cytokines in the respective immune cell subtype from BALFs of COVID-19 patients compared to healthy controls. The edge color denotes the Pearson correlation coefficient, and the edge thickness is proportional to the correlation adjusted P value.

HIF1A (Hypoxia Inducible Factor 1 Alpha) is a master transcriptional regulator that is activated under hypoxic conditions. Indeed, hypoxia is a primary pathophysiological feature in severe COVID-19 and HIF1A is speculated to contribute largely to the CRS by activating and preventing turnover of immune cells including macrophages and neutrophils, which secrete large amounts of inflammatory cytokines (32–35). Consistent with this, we found that in seven immune cell types, the expression of HIF1A was positively correlated with the expression of CCL2, CCL5, and CXCL8, which are potent chemoattractants for immature macrophages and neutrophils (36), and the expression of TNFSF13B, which promotes cell survival (37). These findings suggest that targeting HIF1A could interfere with several processes that contribute to the CRS in COVID-19.

### Targeting TFs to suppress the production of cytokines involved in the COVID-19 CRS

To reduce the expression of cytokines associated with the COVID-19 CRS, we sought drugs that target the major TF hubs within the network. We prioritized drugs by their status as approved or investigational (*i.e*. in clinical trials), selectivity, and availability. Based on these criteria, we selected five FDA approved drugs that target the TF hubs (carvedilol - HIF1A, dexamethasone - NR3C1, dimethyl fumarate - RELA, glycyrrhizic acid - NFKB1/2, and sulfasalazine - NF-κB) and one clinical drug (T5224 - FOS/JUN), and investigated their ability to downregulate several key cytokines implicated in the COVID-19 CRS (CCL2, CXCL8, and IL6).

We investigated the effect of these drugs, alone and in combination, in peripheral blood mononuclear cells (PBMCs) from four healthy human donors stimulated with R848 or LPS, potent TLR7/TLR8 and TLR4 agonists, respectively (38–40). Since TLR7/TLR8 recognize single-stranded RNA from viruses such as SARS-CoV-2 and TLR4, a receptor that recognizes various endogenous and exogenous proteins which was predicted to strongly interact with the SARS-CoV-2 spike glycoprotein (41), activation of these TLR signaling pathways can partially mimic the inflammatory response in COVID-19. We found two drugs, dimethyl fumarate and T5224, that inhibited the production of CCL2, CXCL8, and IL6 (Figure 2C-D). This confirms that targeting TF hubs has the potential to concomitantly limit the production of multiple cytokines upregulated in COVID-19 patients. Additionally, testing all pairwise drug combinations revealed 11 combinations that synergistically reduced the production of at least one cytokine in either stimulated conditions in all PBMC donors (Figure 2C-D). In particular, the combination of dexamethasone with sulfasalazine or T5224 most consistently produced a synergistic effect in reducing cytokine production across the PBMC donors. This may be attributed to these drugs targeting TFs in parallel inflammatory pathways. Collectively, we identified multiple candidate repurposable drugs for the potential treatment of COVID-19 CRS. However, further animal models and clinical trials are required to verify the clinical benefits of these predicted drug candidates.

**Figure 2.**
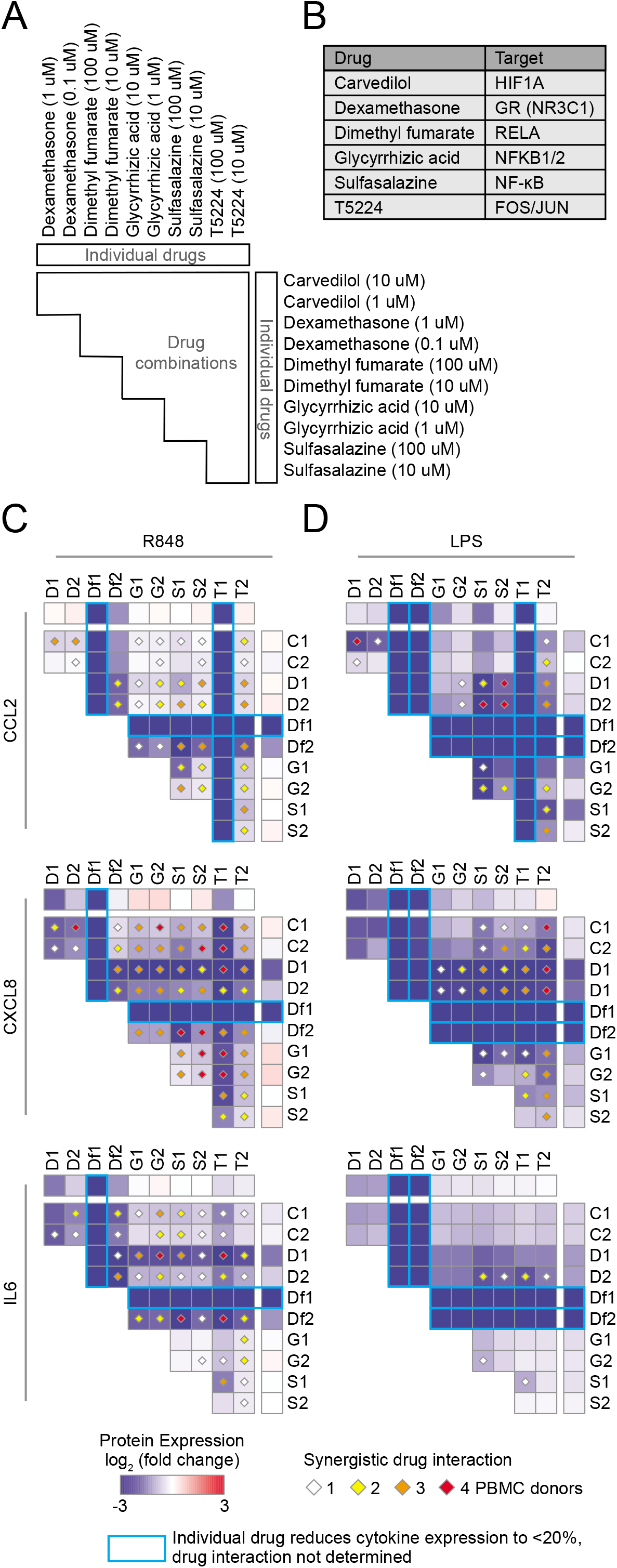
Identification of synergistic drug combinations targeting TF hubs that regulate inflammatory cytokines. (A) Schematic of experimental design to test 60 drug combinations. (B) Drug TF targets. (C-D) Heatmaps showing the average log_2_ (fold change) cytokine production across four PBMC donors treated with the indicated drugs, relative to PBMCs not treated with the indicated drugs, and stimulated with (C) R848 or (D) LPS. Diamonds indicate synergistic drug interactions, as determined by the coefficient of drug interaction, observed in 1 (white), 2 (yellow), 3 (orange), or all 4 (red) PBMC donors. Blue boxes represent cases wherein the individual drug reduced cytokine expression to less than 20%, and therefore synergistic effects were not evaluated.

### Targeting nuclear receptors to suppress the production of cytokines involved in the COVID-19 CRS

TFs from the nuclear receptor (NR) family present promising therapeutic targets because of the lipophilic nature of their ligands and because numerous FDA approved drugs targeting NRs are currently available. Not surprisingly, only a few NRs were significantly correlated in expression with cytokines overexpressed in the COVID-19 patients (Figure 1), since NRs are ligand-activated TFs and therefore their activities are primarily regulated at the protein level. To explore the therapeutic potential of targeting NRs to reduce the expression of cytokines elevated in COVID-19 patients, we first identified cytokines that were significantly upregulated (Padj<0.05, fold change ≥ 2) in the BALFs of moderate (Figure 3A) and severe (Figure 3B) COVID-19 patients compared to healthy controls (12). We then analyzed the expression of these cytokines using publicly available transcriptomic data collected from primary human cells and cell lines treated with small molecule NR drugs (Signaling Pathways Project) (42). We found that, while drugs targeting NRs across many families can modulate the expression of cytokines, drugs targeting members of the 3-ketosteroid, vitamin D, and peroxisome proliferator-activated receptor families, tend to reduce the expression of inflammatory cytokines (Figure 3C).

**Figure 3.**
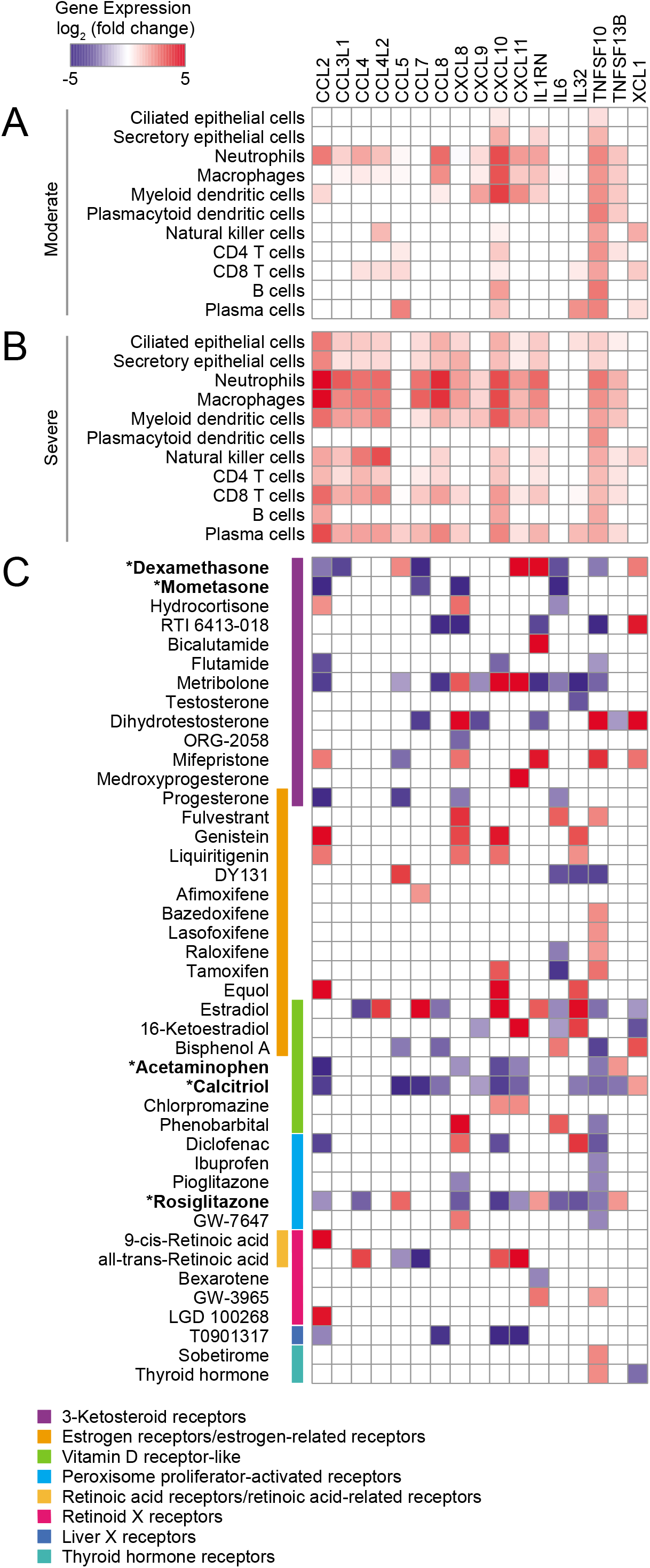
Exploration of repurposable nuclear receptor drugs. (A-B) Heatmaps showing the average log_2_ (fold change) cytokine gene expression in the indicated cell types from BALFs of (A) moderate and (B) severe COVID-19 patients relative to healthy controls. (C) Heatmap showing the average log_2_ (fold change) cytokine gene expression in response to treatment with small molecule NR drugs. Data was obtained from the Signaling Pathways Project Transcriptomine resource.

We next investigated the therapeutic potential of five approved NR drugs (acetaminophen, dexamethasone, ercalcitriol, mometasone, and rosiglitazone) that strongly downregulated the expression of multiple cytokines in the expression profiling datasets (Figure 3C), and assayed their effect on CCL2, CXCL8, and IL6, in R848 or LPS stimulated PBMCs (Figure 4A-D). Indeed, TF targets of some of these drugs, for example NR3C1 and VDR, have been reported to directly regulate CCL2, CXCL8, and IL6 expression in other stimulated contexts (9, 10, 43). We found that dexamethasone and mometasone, both of which target NR3C1, most potently reduced the production of CXCL8 and IL6 (Figure 4C-D). Additionally, testing all pairwise combinations revealed that all 10 drug combinations either additively or synergistically reduced the production of CCL2 and CXCL8 in all PBMC samples stimulated with R848 (Figure 4C). Generally, across the PBMC samples, combinations of dexamethasone with rosiglitazone and mometasone with ercalcitriol most consistently produced a synergistic effect in reducing cytokine production, while combinations of dexamethasone or mometasone with ercalcitriol most potently reduced cytokine production. Overall, these findings suggest there may be potential therapeutic benefits of repurposing these NR drugs to suppress the CRS in COVID-19 patients. Further studies are required to determine the clinical benefits of these drug candidates.

**Figure 4.**
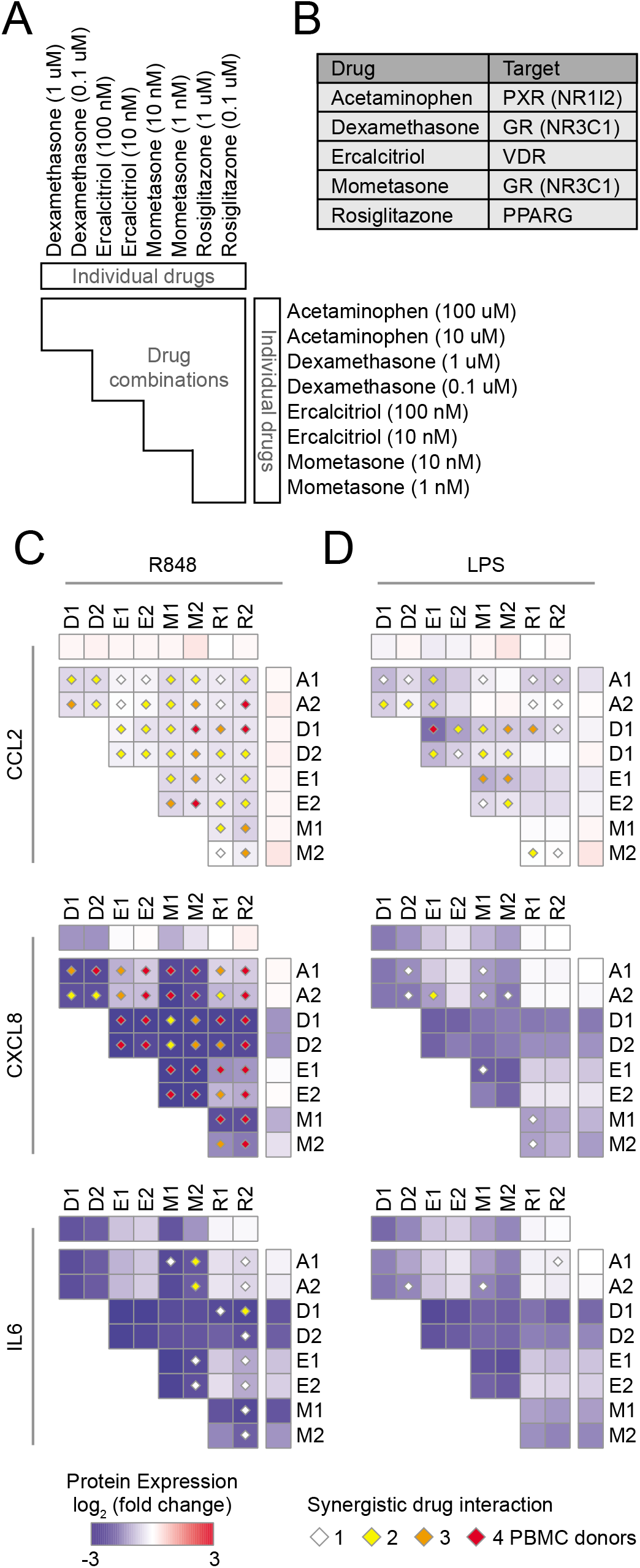
Identification of synergistic drug combinations targeting nuclear receptors that regulate inflammatory cytokines. (A) Schematic of experimental design to test 40 drug combinations. (B) Drug NR TF targets. (C-D) Heatmaps showing the average log_2_ (fold change) cytokine production across four PBMC donors treated with the indicated drugs, relative to PBMCs not treated with the indicated drugs, and stimulated with (C) R848 or (D) LPS. Diamonds indicate synergistic drug interactions, as determined by the coefficient of drug interaction, observed in 1 (white), 2 (yellow), 3 (orange), or all 4 (red) PBMC donors.

## Discussion

In the present study, we used a gene regulatory network approach to identify candidate TFs that regulate cytokines overexpressed in COVID-19 patients and evaluated approved drugs targeting these TFs for their ability to downregulate three key cytokines frequently associated with disease severity. We identified two drugs (dimethyl fumarate and T5224) that individually potently suppressed cytokine production, and 25 drug combinations that could synergistically suppress cytokine production. Altogether, these findings provide several promising candidate drugs and targets with potential therapeutic effects for controlling the CRS in COVID-19.

Our network-based approach identified TF hubs that likely regulate many of the cytokines overexpressed in COVID-19 patients. We showed that by targeting these TF hubs, for example targeting RELA with dimethyl fumarate and AP-1 with T5224, we were able to concomitantly inhibit the production of multiple cytokines. Moreover, targeting TF hubs may also interfere with positive feedback and feedforward loops of cytokine production that lead to the CRS (23). It would also be interesting to explore targeting non-hub TFs that regulate a key cytokine responsible for driving these loops, as the effects could me more specific with less side effects.

Combination therapies have the potential to increase drug efficacy and reduce side effects, and have thus become a routine strategy in the treatment of diseases (44). In particular, synergistic combinations allow the use of lower doses to achieve the same effect as the individual drugs, which may reduce adverse reactions. Notably, nearly all drugs we tested achieved a similar or stronger suppression of cytokine production when used at a 10-fold lower dose in combination than when used individually. This includes the combination of dexamethasone with ercalcitriol (active metabolite of vitamin D). Thus, in the debate of whether vitamin D supplementation has beneficial effects in the treatment of COVID-19, at least from the perspective of treating the CRS, vitamin D may enhance the anti-inflammatory effects of dexamethasone.

Aside from having well-known anti-inflammatory properties, some of the drugs tested also have reported antiviral properties against SARS-CoV-2. Dimethyl fumarate, mometasone, calcitriol, and sulfasalazine, potently inhibited SARS-CoV-2 replication *in vitro* in Vero E6 cells (45–48). The antiviral activities of Glycyrrhizin have been extensively studied in the context of other human viruses and most notably, the drug was found to potently suppress replication of two clinical isolates of SARS-associated coronavirus in Vero cells (49), and preliminarily, neutralize SARS-CoV-2 by inhibiting the viral main protease (50). Further, Carvedilol and Acetaminophen have been reported to decrease the expression of ACE2 and serine protease TMPRSS2, respectively (51, 52), both of which are required for SARS-CoV-2 entry into cells (53). Hence, drug combinations that simultaneously exert both anti-inflammatory and antiviral effects against SARS-CoV-2 may have the greatest potential to be effective in treating COVID-19.

There is growing evidence that certain food supplements may have therapeutic benefits in COVID-19. Numerous small-scale studies have found that patients with sufficient vitamin D levels are less likely to have life-threatening complications (54–56). Additionally, glycyrrhizic acid, a frequent component in traditional Chinese medicines and the main constituent in licorice, has been reported to have anti-inflammatory properties, by antagonizing TLR4 (57, 58), and broad antiviral activities (49). Other foods are also known to inhibit inflammatory mediators, for example curcumin, a substance in turmeric that gives curry its distinct flavor and yellow color, inhibits numerous TFs including NF-κB, AP-1, and HIF1A, and has potent anti-inflammatory properties (59, 60). Hence, a study of the association between food intake and the severity of COVID-19 symptoms and outcomes may shed light into differences in severity and mortality between countries (61).

In summary, our approach of targeting transcriptional regulators of cytokines associated with the CRS provides candidate drugs and targets to treat COVID-19. However, additional research is needed to determine whether these combinations elicit the same immunomodulatory response in the context of SARS-CoV-2 infection. More importantly, although all the drugs investigated in this study are FDA approved, careful evaluation of the efficacy, safety, and risk-benefit balance of these drugs in animal models and COVID-19 patients is necessary as outcomes of drug interactions could drastically differ between *in vitro, in vivo*, and clinical trials. Nonetheless, the candidate drugs show promise for further investigation in downregulating the CRS in COVID-19 patients. More broadly, the findings reported here may also be applicable to CRS resulting from other viral infections, bacterial infections, sepsis, and CAR-T therapies.

## Methods

### ScRNA-seq data processing

scRNA-seq data from BALF cells of COVID-19 patients and healthy controls was downloaded from GEO repositories GSE145926 (12) and GSE128033 (15). For all datasets, we used STARsolo (v 2.7.3) (62) to align reads to the human GRCh38 genome and quantify read counts to determine gene expression. We used Scrubblet (63) to detect and remove doublets, and then filtered the remaining data to only retain cells with 1000-50000 UMI counts, 500-7500 genes, and less than 25% mitochondrial reads. A total of 72,433 cells remaining were used for all subsequent analysis. Finally, the data was normalized using the regularized negative binomial regression method (64) and batch effect was removed using the Canonical Correlation Analysis method (65).

Cell clustering was performed using Seurat (v3.1.4) (65) and cell type classifications were obtained using SingleR (66), and then validated with canonical immune cell type markers. The following markers were used to identify cell types: ciliated epithelial cells: TUBB4B and TPPP3; secretory epithelial cells: SCGB3A1 and SCGB1A1; neutrophils: S100A8, S100A9 and FCN1; macrophages: APOE, C1QA and C1QB; myeloid dendritic cells: FCER1A and CD1C; plasmacytoid dendritic cells: TCF4 and TNFRSF21; mast cells: AREG, TPSB2 and TPSAB1; NK cells: GNLY, PRF1, NKG7 and the absence of the general T cell markers; T cells: CD3D, CD3G, CD4E, CD4 (CD4 T cells only) and CD8 (CD8 T cells only); B cells: CD79A, CD79B and MS4A1; plasma cells: IGHG1, IGHG2 and IGHG4.

### Differential gene expression analysis

Differential gene expression analysis, between BALF cells from COVID-19 patients and healthy controls, was performed using a Wilcoxon test, and the P values were adjusted by false discovery rate correction using the Benjamini-Hochberg method.

### Correlation analysis

Correlation coefficients between TFs and cytokines in BALF cells from COVID-19 patients were determined using the Pearson correlation method, and the P values were adjusted by false discovery rate correction using the Benjamini-Hochberg method. The correlation analyses were restricted to cells with reads for both the TF and the cytokine, to limit noise and over-estimation of the correlation due to cells with zero reads for either the TF or the cytokine or both, and cell types with more than 10% of cells expressing both the TF and the cytokine. Correlations between TFs and cytokines were determined per cell type.

### Signaling Pathways Project data acquisition and processing

A list of cytokines that were differentially expressed and upregulated in BALFs of COVID-19 patients compared to healthy controls was submitted to the Signaling Pathways Project Ominer web tool (42) on July 25, 2020. The search criteria included Omics Category: Transcriptomics, Module Category: Nuclear receptors - all families, Biosample Category: Human - all physiological systems, FDR Significance Cut-off: 5E-02. The search results were downloaded as a table reporting the fold change for cytokine gene expression in experimental versus control conditions. Only experiments involving small molecule NR drugs were further explored in our analysis. If there were multiple experiments for a drug-cytokine interaction, only interactions wherein at least 80% of the experiments resulted in cytokine gene expression changing in the same direction were included in our analysis. Finally, for each drug-cytokine interaction, we calculated the median fold change in cytokine gene expression across the experiments and depicted the data in a heatmap.

### PBMC purification and drug treatment

Peripheral blood mononuclear cells (PBMCs) were isolated from de-identified human leukapheresis-processed blood (New York Biologics, Inc) by centrifugation through Lymphoprep (Stem Cell Technologies.) density gradient. PBMCs were washed in PBS, resuspended in red blood cell lysis solution for 5 min, and washed three more times in PBS. Purified PBMCs were cultured in RPMI supplemented with 10% FBS and 1% Antibiotic-Antimycotic (100X) and plated in 96-well plates at a density of 1 × 10^6^ cells/ml and 0.1 ml/well. Purified PBMCs were immediately treated with the different drugs or frozen in RPMI supplemented with 40% FBS, 10% DMSO, and 1% Antibiotic-Antimycotic (100X). Frozen PBMCs were rapidly thawed in a 37°C water bath, washed three times in warm RPMI supplemented with 10% FBS and 1% Antibiotic-Antimycotic (100X), and rested for 1 hour at 37°C before drug treatment. Fresh or thawed PBMCs were pretreated with Acetaminophen (MiliporeSigma), Dexamethasone (MiliporeSigma), Ercalcitriol (Tocris), Mometasone (Tocris), or Rosiglitazone (Tocris), at the various concentrations for 30 minutes, and then stimulated with R848 (1 μM) or LPS (100 ng/ml), for 20 hours. The supernatants were collected and the amounts of CCL2, CXCL8, and IL6, were quantified by ELISA. Each experimental condition was performed in two biological replicates for each PBMC donor, and the average of the replicates was used to determine cytokine expression.

### Measurement of cytokine production

The amount of cytokines (CCL2, CXCL8, and Il6) in treated PBMC supernatants were quantified by ELISA using the ELISA MAX Deluxe Set Human CCL2 (Biolegend), ELISA MAX Deluxe Set Human IL8 (Biolegend), and ELISA MAX Deluxe Set Human IL6 (Biolegend) kits according to the manufacturer’s protocol.

### Calculation of coefficient of drug interaction

To determine drug interactions (*i.e*. additive, synergistic, or antagonistic), we calculated the coefficient of drug interaction (CDI) using the formula CDI=AB/(A×B), where AB is the ratio of the combination to the control, and A or B is the ratio of the single drug to the control. We then applied the following thresholds to determine the drug interaction, CDI=0.7-1.3 indicates an additive interaction, CDI <0.7 indicates a synergistic interaction, and CDI >1.3 indicates an antagonistic interaction.

## Acknowledgements

This work was funded by the National Institutes of Health grant R35 GM128625 awarded to J.I.F.B.

## Author contributions

J.I.F.B. conceived and supervised the project. C.S.S. designed the experiments, and C.S.S. and X.L. performed the experiments. C.S.S., L.Z., J.T.R., and V.X.S. performed the data analysis. C.S.S., V.X.S., and J.I.F.B. wrote the manuscript. All authors read and approved the manuscript.

---

**Supplementary Figure 1. scRNA-seq data of BALF cells from COVID-19 patients and healthy subjects.**

(A) Uniform Manifold Approximation and Projection (UMAP) plots presenting clusters and color-coded major cell types identified in single cell transcriptomes of BALF cells from moderate and severe COVID-19 (n = 9) patients and healthy (n = 3) subjects. (B) The average expression and percentage of expression of cell calling markers in each cell population.

**Supplementary Table 1.** List of drugs for TFs in the COVID-19 cytokine gene regulatory networks.

## References

1. Dong E, Du H, & Gardner L (2020) An interactive web-based dashboard to track COVID-19 in real time. Lancet Infect Dis 20(5):533–534.

2. Gandhi RT, Lynch JB, & Del Rio C (2020) Mild or Moderate Covid-19. N Engl J Med 383(18):1757–1766.

3. Mehta P, et al. (2020) COVID-19: consider cytokine storm syndromes and immunosuppression. Lancet 395(10229):1033–1034.

4. Kalil AC, et al. (2020) Baricitinib plus Remdesivir for Hospitalized Adults with Covid-19. N Engl J Med.

5. Group RC, et al. (2020) Dexamethasone in Hospitalized Patients with Covid-19 - Preliminary Report. N Engl J Med.

6. Beigel JH, et al. (2020) Remdesivir for the Treatment of Covid-19 - Final Report. N Engl J Med 383(19):1813–1826.

7. Cronstein BN, Kimmel SC, Levin RI, Martiniuk F, & Weissmann G (1992) A mechanism for the antiinflammatory effects of corticosteroids: the glucocorticoid receptor regulates leukocyte adhesion to endothelial cells and expression of endothelial-leukocyte adhesion molecule 1 and intercellular adhesion molecule 1. Proc Natl Acad Sci U S A 89(21):9991–9995.

8. Lee DW, et al. (2014) Current concepts in the diagnosis and management of cytokine release syndrome. Blood 124(2):188–195.

9. Sasse SK, et al. (2016) Glucocorticoid and TNF signaling converge at A20 (TNFAIP3) to repress airway smooth muscle cytokine expression. Am J Physiol Lung Cell Mol Physiol 311(2):L421–432.

10. Kadiyala V, et al. (2016) Cistrome-based Cooperation between Airway Epithelial Glucocorticoid Receptor and NF-kappaB Orchestrates Anti-inflammatory Effects. J Biol Chem 291(24): 12673–12687.

11. Wishart DS, et al. (2018) DrugBank 5.0: a major update to the DrugBank database for 2018. Nucleic Acids Res 46(D1):D1074–D1082.

12. Liao M, et al. (2020) Single-cell landscape of bronchoalveolar immune cells in patients with COVID-19. Nat Med 26(6):842–844.

13. Zhou Z, et al. (2020) Heightened Innate Immune Responses in the Respiratory Tract of COVID-19 Patients. Cell Host Microbe 27(6):883–890 e882.

14. Santoso CS, et al. (2020) Comprehensive mapping of the human cytokine gene regulatory network. Nucleic Acids Res 48(21):12055–12073.

15. Morse C, et al. (2019) Proliferating SPP1/MERTK-expressing macrophages in idiopathic pulmonary fibrosis. Eur Respir J 54(2).

16. Tamura T, Yanai H, Savitsky D, & Taniguchi T (2008) The IRF family transcription factors in immunity and oncogenesis. Annu Rev Immunol 26:535–584.

17. Park A & Iwasaki A (2020) Type I and Type III Interferons - Induction, Signaling, Evasion, and Application to Combat COVID-19. Cell Host Microbe 27(6):870–878.

18. Zhang Q, et al. (2020) Inborn errors of type I IFN immunity in patients with life-threatening COVID-19. Science 370(6515).

19. Bastard P, et al. (2020) Autoantibodies against type I IFNs in patients with life-threatening COVID-19. Science 370(6515).

20. Lee JS & Shin EC (2020) The type I interferon response in COVID-19: implications for treatment. Nat Rev Immunol 20(10):585–586.

21. Israelow B, et al. (2020) Mouse model of SARS-CoV-2 reveals inflammatory role of type I interferon signaling. J Exp Med 217(12).

22. Stebbing J, et al. (2020) JAK inhibition reduces SARS-CoV-2 liver infectivity and modulates inflammatory responses to reduce morbidity and mortality. Sci Adv.

23. Hirano T & Murakami M (2020) COVID-19: A New Virus, but a Familiar Receptor and Cytokine Release Syndrome. Immunity 52(5):731–733.

24. Martin T, Cardarelli PM, Parry GC, Felts KA, & Cobb RR (1997) Cytokine induction of monocyte chemoattractant protein-1 gene expression in human endothelial cells depends on the cooperative action of NF-kappa B and AP-1. Eur J Immunol 27(5):1091–1097.

25. Akira S, Taga T, & Kishimoto T (1993) Interleukin-6 in biology and medicine. Adv Immunol 54:1–78.

26. Huang C, et al. (2020) Clinical features of patients infected with 2019 novel coronavirus in Wuhan, China. Lancet 395(10223):497–506.

27. Merad M & Martin JC (2020) Pathological inflammation in patients with COVID-19: a key role for monocytes and macrophages. Nat Rev Immunol 20(6):355–362.

28. Blanco-Melo D, et al. (2020) Imbalanced Host Response to SARS-CoV-2 Drives Development of COVID-19. Cell 181(5):1036–1045 e1039.

29. Satoh T, et al. (1988) Induction of neuronal differentiation in PC12 cells by B-cell stimulatory factor 2/interleukin 6. Mol Cell Biol 8(8):3546–3549.

30. Lord KA, Abdollahi A, Hoffman-Liebermann B, & Liebermann DA (1993) Proto-oncogenes of the fos/jun family of transcription factors are positive regulators of myeloid differentiation. Mol Cell Biol 13(2):841–851.

31. Lin YM, Hsu CJ, Liao YY, Chou MC, & Tang CH (2012) The CCL2/CCR2 axis enhances vascular cell adhesion molecule-1 expression in human synovial fibroblasts. PLoS One 7(11):e49999.

32. Walmsley SR, et al. (2005) Hypoxia-induced neutrophil survival is mediated by HIF-1alpha-dependent NF-kappaB activity. J Exp Med 201(1):105–115.

33. Nizet V & Johnson RS (2009) Interdependence of hypoxic and innate immune responses. Nat Rev Immunol 9(9):609–617.

34. Marchetti M (2020) COVID-19-driven endothelial damage: complement, HIF-1, and ABL2 are potential pathways of damage and targets for cure. Ann Hematol 99(8):1701–1707.

35. Jahani M, Dokaneheifard S, & Mansouri K (2020) Hypoxia: A key feature of COVID-19 launching activation of HIF-1 and cytokine storm. J Inflamm (Lond) 17(1):33.

36. Sokol CL & Luster AD (2015) The chemokine system in innate immunity. Cold Spring Harb Perspect Biol 7(5).

37. Lee JW, Lee J, Um SH, & Moon EY (2017) Synovial cell death is regulated by TNF-alpha-induced expression of B-cell activating factor through an ERK-dependent increase in hypoxia-inducible factor-1alpha. Cell Death Dis 8(4):e2727.

38. Jurk M, et al. (2002) Human TLR7 or TLR8 independently confer responsiveness to the antiviral compound R-848. Nat Immunol 3(6):499.

39. Hemmi H, et al. (2002) Small anti-viral compounds activate immune cells via the TLR7 MyD88-dependent signaling pathway. Nat Immunol 3(2):196–200.

40. Chow JC, Young DW, Golenbock DT, Christ WJ, & Gusovsky F (1999) Toll-like receptor-4 mediates lipopolysaccharide-induced signal transduction. J Biol Chem 274(16):10689–10692.

41. Choudhury A & Mukherjee S (2020) In silico studies on the comparative characterization of the interactions of SARS-CoV-2 spike glycoprotein with ACE-2 receptor homologs and human TLRs. J Med Virol 92(10):2105–2113.

42. Ochsner SA, et al. (2019) The Signaling Pathways Project, an integrated ‘omics knowledgebase for mammalian cellular signaling pathways. Sci Data 6(1):252.

43. Masood R, et al. (2000) Kaposi sarcoma is a therapeutic target for vitamin D(3) receptor agonist. Blood 96(9):3188–3194.

44. Sun W, Sanderson PE, & Zheng W (2016) Drug combination therapy increases successful drug repositioning. Drug Discov Today 21(7):1189–1195.

45. Olagnier D, et al. (2020) SARS-CoV2-mediated suppression of NRF2-signaling reveals potent antiviral and anti-inflammatory activity of 4-octyl-itaconate and dimethyl fumarate. Nat Commun 11(1):4938.

46. Matsuyama S, et al. (2020) The Inhaled Steroid Ciclesonide Blocks SARS-CoV-2 RNA Replication by Targeting the Viral Replication-Transcription Complex in Cultured Cells. J Virol 95(1).

47. Chee Keng Mok YLN, Bintou Ahmadou Ahidjo, Regina Ching Hua Lee, Marcus Wing Choy Loe, Jing Liu, Kai Sen Tan, Parveen Kaur, Wee Joo Chng, John Eu-Li Wong, De Yun Wang, Erwei Hao, Xiaotao Hou, Yong Wah Tan, Tze Minn Mak, Cui Lin, Raymond Lin, Paul Tambyah, JiaGang Deng, Justin Jang Hann Chu (2020) Calcitriol, the active form of vitamin D, is a promising candidate for COVID-19 prophylaxis. BioRxiv (https://doi.org/10.1101/2020.06.21.162396).

48. Namshik Han WH, Konstantinos Tzelepis, Patrick Schmerer, Eliza Yankova, Méabh MacMahon, Winnie Lei, Nicholas M Katritsis, Anika Liu, Alison Schuldt, Rebecca Harris, Kathryn Chapman, Frank McCaughan, Friedemann Weber, Tony Kouzarides (2020) Identification of SARS-CoV-2 induced pathways reveal drug repurposing strategies. BioRxiv: https://doi.org/10.1101/2020.1108.1124.265496.

49. Cinatl J, et al. (2003) Glycyrrhizin, an active component of liquorice roots, and replication of SARS-associated coronavirus. Lancet 361(9374):2045–2046.

50. L. van de Sand MB, M. Alt, L. Schipper, C.S. Heilingloh, D. Todt, U. Dittmer, C. Elsner, O. Witzke, View ORCID ProfileA. Krawczyk (2020) Glycyrrhizin effectively neutralizes SARS-CoV-2 in vitro by inhibiting the viral main protease. BioRxiv: https://doi.org/10.1101/2020.1112.1118.423104.

51. Skayem C & Ayoub N (2020) Carvedilol and COVID-19: A Potential Role in Reducing Infectivity and Infection Severity of SARS-CoV-2. Am J Med Sci 360(3):300.

52. Aleksei Zarubin VS, Anton Markov, Nikita Kolesnikov, Andrey Marusin, Irina Khitrinskaya, Maria Swarovskaya, Sergey Litvinov, Natalia Ekomasova, Murat Dzhaubermezov, Nadezhda Maksimova, Aitalina Sukhomyasova, Olga Shtygasheva, Elza Khusnutdinova, Magomed Radjabov, Vladimir Kharkov (2020) Structural variability, expression profile and pharmacogenetics properties of TMPRSS2 gene as a potential target for COVID-19 therapy. BioRxiv: https://doi.org/10.1101/2020.1106.1120.156224.

53. Hoffmann M, et al. (2020) SARS-CoV-2 Cell Entry Depends on ACE2 and TMPRSS2 and Is Blocked by a Clinically Proven Protease Inhibitor. Cell 181(2):271–280 e278.

54. Meltzer DO, et al. (2020) Association of Vitamin D Status and Other Clinical Characteristics With COVID-19 Test Results. JAMA Netw Open 3(9):e2019722.

55. Jain A, et al. (2020) Analysis of vitamin D level among asymptomatic and critically ill COVID-19 patients and its correlation with inflammatory markers. Sci Rep 10(1):20191.

56. Grant WB, et al. (2020) Evidence that Vitamin D Supplementation Could Reduce Risk of Influenza and COVID-19 Infections and Deaths. Nutrients 12(4).

57. Murck H (2020) Symptomatic Protective Action of Glycyrrhizin (Licorice) in COVID-19 Infection? Front Immunol 11:1239.

58. Bailly C & Vergoten G (2020) Glycyrrhizin: An alternative drug for the treatment of COVID-19 infection and the associated respiratory syndrome? Pharmacol Ther 214:107618.

59. Singh S & Aggarwal BB (1995) Activation of transcription factor NF-kappa B is suppressed by curcumin (diferuloylmethane) [corrected]. J Biol Chem 270(42):24995–25000.

60. Bae MK, et al. (2006) Curcumin inhibits hypoxia-induced angiogenesis via down-regulation of HIF-1. Oncol Rep 15(6):1557–1562.

61. Samaddar A, Gadepalli R, Nag VL, & Misra S (2020) The Enigma of Low COVID-19 Fatality Rate in India. Front Genet 11:854.

62. Dobin A, et al. (2013) STAR: ultrafast universal RNA-seq aligner. Bioinformatics 29(1):15–21.

63. Wolock SL, Lopez R, & Klein AM (2019) Scrublet: Computational Identification of Cell Doublets in Single-Cell Transcriptomic Data. Cell Syst 8(4):281–291 e289.

64. Hafemeister C & Satija R (2019) Normalization and variance stabilization of single-cell RNA-seq data using regularized negative binomial regression. Genome Biol 20(1):296.

65. Butler A, Hoffman P, Smibert P, Papalexi E, & Satija R (2018) Integrating single-cell transcriptomic data across different conditions, technologies, and species. Nat Biotechnol 36(5):411–420.

66. Aran D, et al. (2019) Reference-based analysis of lung single-cell sequencing reveals a transitional profibrotic macrophage. Nat Immunol 20(2):163–172.

