## Supplementary figures and images for "*In vitro* Targeting of Transcription Factors to Control the Cytokine Release Syndrome in COVID-19"

### Supplemental Figure 1

Supplementary Figure 1

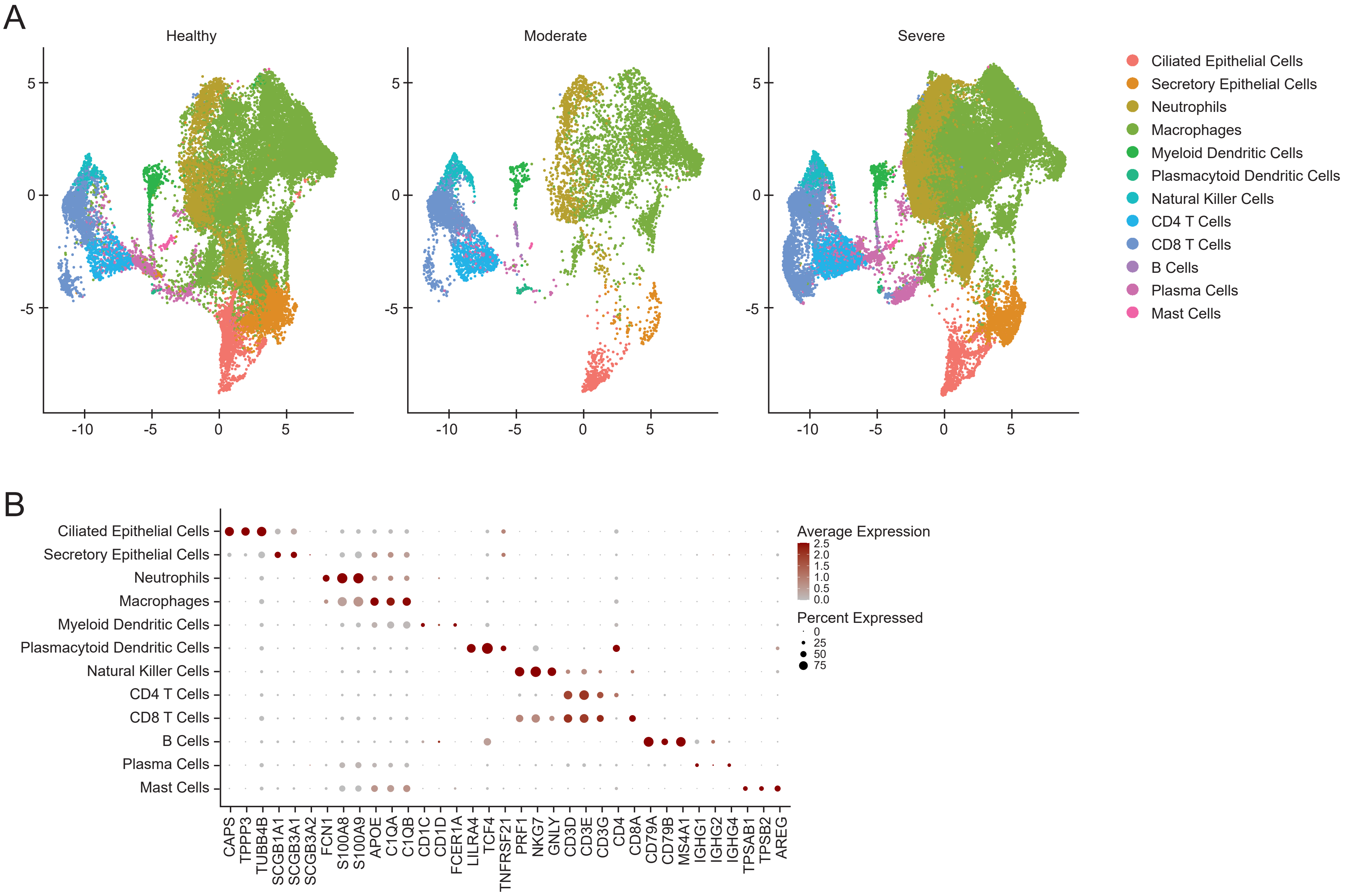
