## Supplemental Table 1 for "*In vitro* Targeting of Transcription Factors to Control the Cytokine Release Syndrome in COVID-19"

| TARGET | DRUGS |
|---|---|
| **AHR** | Atorvastatin, Diosmin, Flutamide, Ginseng, Leflunomide, Omeprazole |
| **ATF2** | Pseudoephedrine |
| **ATF3** | Pseudoephedrine |
| **ATF4** | Pseudoephedrine |
| **ATF7** | Pseudoephedrine |
| **CREB1** | Adenosine phosphate, Naloxone |
| **FOS** | Nadroparin, Pseudoephedrine, T5224 |
| **HIF1A** | Carvedilol, Hydralazine |
| **JUN** | Irbesartan, Pseudoephedrine, T5224 |
| **MYC** | Acetylsalicylic acid (Aspirin), Nadroparin, |
| **NFKB1** | Donepezil, Glycyrrhizic acid, Pseudoephedrine, Triflusal |
| **NFKB2** | Donepezil, Glucosamine, Glycyrrhizic acid, |
| **NR3C1** | Betamethasone, Budesonide, Dexamethasone, Fluticasone, Hydrocortisone, Mometasone, Methylprednisolone, Prednisolone, Prednisone, Triamcinolone |
| **NR4A3** | Dasatinib |
| **RELA** | Dimethyl fumarate |
| **RORA** | Cholesterol |
| **RXRA** | Acitretin, Alitretinoin, Alpha-Linolenic acid, Doconexent, Etodolac, Isotretinoin, Oleic acid, Rosiglitazone |
| **STAT5B** | Dasatinib |
| **THRB** | Levothyroxine, Liothyronine, Liotrix |

*Data collected from DrugBank (https://go.drugbank.com/)
